# Multivariate Meta-Analysis of Differential Principal Components underlying Human Primed and Naive-like Pluripotent States

**DOI:** 10.1101/2020.10.20.347666

**Authors:** Kory R. Johnson, Barbara S. Mallon, Yang C. Fann, Kevin G. Chen

**Author notes:** **Correspondence to**: Dr. Kory R. Johnson, Dr. Kevin G. Chen.

## Abstract

The ground or naive pluripotent state of human pluripotent stem cells (hPSCs), which was initially established in mouse embryonic stem cells (mESCs), is an emerging and tentative concept. To verify this important concept in hPSCs, we performed a multivariate meta-analysis of major hPSC datasets via the combined analytic powers of percentile normalization, principal component analysis (PCA), *t*-distributed stochastic neighbor embedding (*t*-SNE), and SC3 consensus clustering. This vigorous bioinformatics approach has significantly improved the predictive values of the current meta-analysis. Accordingly, we were able to reveal various similarities between some naive-like hPSCs (NLPs) and their human and mouse *in vitro* counterparts. Moreover, we also showed numerous fundamental inconsistencies between diverse naive-like states, which are likely attributed to interlaboratory protocol differences. Collectively, our meta-analysis failed to provide global transcriptomic markers that support a bona fide human naive pluripotent state, rather suggesting the existence of altered pluripotent states under current naive-like growth protocols.

---

Ground or naive states in pluripotent stem cells were initially proposed by Smith and colleagues based on the identification of lineage-primed epiblast stem cells (EpiSCs) derived from post-implantation mouse embryos (Brons et al., 2007; Tesar et al., 2007) and on the studies of naive mouse embryonic stem cells (mESCs) from pre-implantation mouse embryos (Nichols and Smith, 2009; Ying et al., 2008). Hence, mouse pluripotent stem cells have two distinct (primed and naive) states. The maintenance of the naive state relies on the use of leukemia inhibitory factor (LIF) with two inhibitors, GSK-3βi and ERK1/2i (abbreviated as 2i), which suppress glycogen synthase kinase-3β (GSK-3β) and extracellular signal-regulated kinases 1/2 (ERK1/2), respectively. Conceivably, pluripotent stem cells in the naive state may have several potential advantages over the primed state, particularly for facilitating single-cell growth, genetic manipulation, disease-modeling, and drug discovery [reviewed in references (Chen et al., 2014; Hanna et al., 2010)].

In the past seven years, several groups have reported the conversion of primed human pluripotent stem cells (hPSCs), which depend on distinct growth signals that embrace FGF2/Activin-A/TGFβ signaling pathways, to naive-like hPSCs (NLPs) and *de novo* derivation of NLPs from the human inner cell mass (Chan et al., 2013; Gafni et al., 2013; Guo et al., 2016; Liu et al., 2017; Takashima et al., 2014; Theunissen et al., 2016; Valamehr et al., 2014; Ware et al., 2014). However, there is a lack of robust assays that precisely define a naive pluripotent state under different growth conditions *in vitro*. The existing assays used for defining pluripotent and differentiation states largely count on various genome-wide analyses (Bock et al., 2011; Chan et al., 2013; Gafni et al., 2013; Takashima et al., 2014; Theunissen et al., 2016). However, genome-wide transcriptomic levels across datasets generated from different laboratories using different technologies (e.g., microarray and RNA-sequencing) often have substantial differences in expression scale and spread. Direct meta-analysis of the transcriptomic levels across datasets can render confusing results and lead to incorrect interpretations and conclusions. Accordingly, previous genome-wide data analyses revealed significant differences between various NLPs derived from different laboratory protocols (Takashima et al., 2014; Theunissen et al., 2016), hence confounding the definition of human naive pluripotency. Thus, there is a pressing need to address the above critical issues.

In this study, we employed a meta-analysis approach that integrates genome-wide microarrays and RNA sequencing (RNA-seq) data into the principal component analysis (PCA) (Jolliffe, 2002), *t*-distributed stochastic neighbor embedding (*t*-SNE) (van der Maaten and Hinton, 2008), and SC3 consensus clustering (Kiselev et al., 2017). We aim to resolve critical interlaboratory experimental inconsistencies with respect to human naive pluripotency. With this approach, we characterized transcriptomic signatures of NLPs from publicly available datasets, and systematically evaluated data from current human naive-like protocols. Our analysis revealed the existence of various naive-like pluripotent states in converted and derived NLPs, which are deficient in global transcriptomic signatures of naive pluripotency as described in both mESCs and early human embryos. Our study also provides new insights into the role of 1D- and 2D-meta-analysis in gene cluster rearrangements, thereby helping define accurate pluripotent states.

## RESULTS

### Assembly of a meta-analysis platform for differentiating pluripotent stem cell states

We have established a multivariate meta-analysis by integrating a rigorously normalized data matrix into PCA, *t-*SNE, and SC3 consensus clustering (Fig. 1). This meta-analysis platform consists of 10 independent modules (M1-10) that build up three major components, which include a gene expression matrix input, three interrelated analytic tools (i.e., PCA, *t*-SNE, and SC3 consensus clustering), and SC3 cluster integration (into PCA and t-SNE) with gene marker discovery modules (Fig. 1). One of the biggest challenges of this meta-analysis was to integrate RNA-seq datasets (n = 6) with microarray datasets (n = 7) for a comparative analysis (Supplementary Tables 1-4). Beside a routine quantile normalization, we utilized an additional percentile normalization followed by a percentile noise exclusion step. This integrated approach only allows the comparison between datasets with mutually expressed genes greater than the 25^th^ percentile. It also significantly increases the comparability between RNA-seq and microarray datasets (Supplementary Fig. 1).

**Figure 1.**
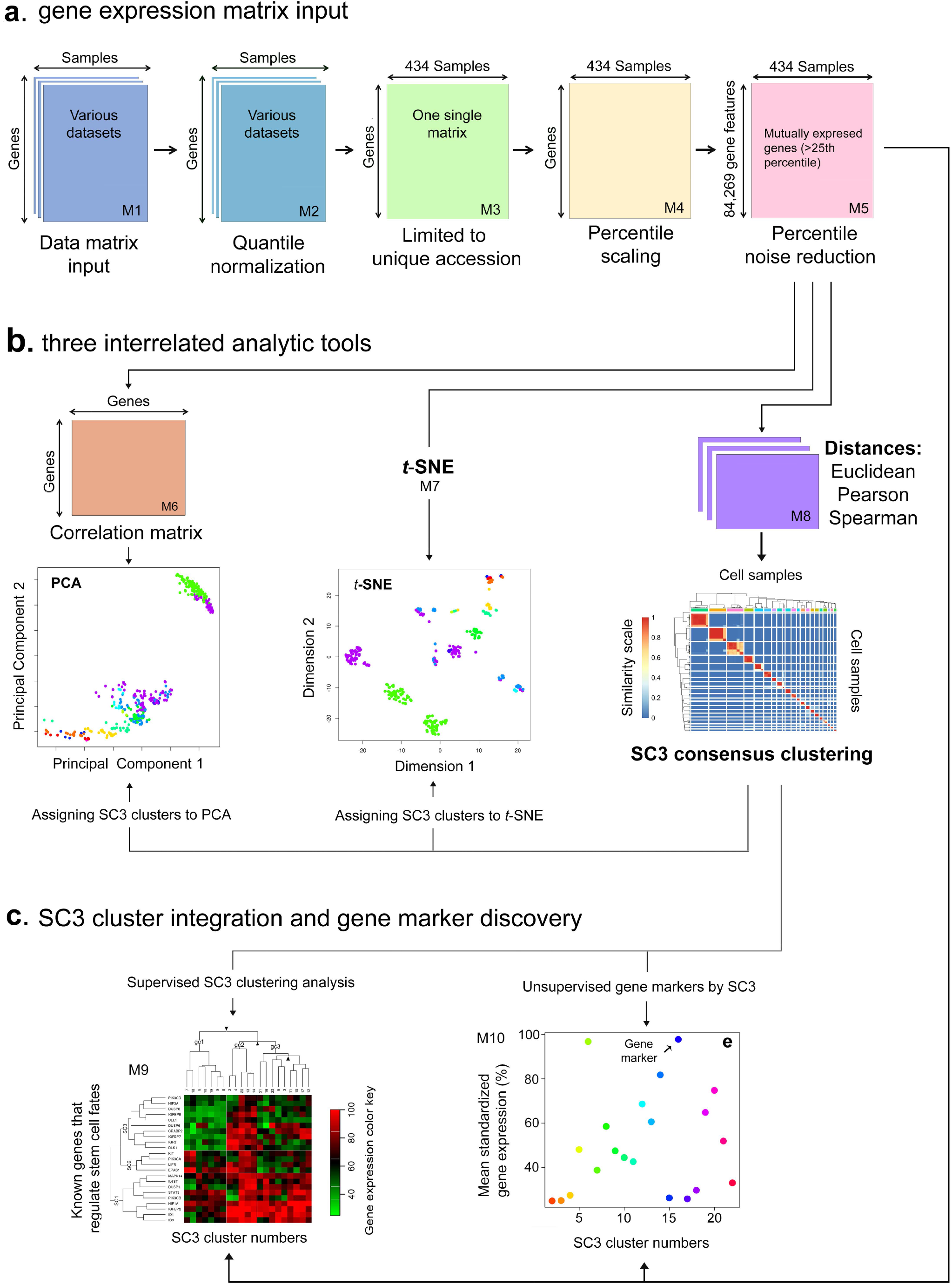
Scheme of a multivariate meta-analysis. The meta-analysis platform consists 10 independent modules (M1-10) that constitute three major components, which include (**a**) gene expression matrix input, that consists 13 datasets and 434 cell samples, classifies cellular states, and employs a 4-step transformation (i.e., quantile normalization, limited to unique genes, collapsing median values, and percentile scaling followed by the exclusion of mRNA noise. (**b**) three interrelated analytic tools (i.e., correlation PCA, *t*-SNE, and SC3 consensus clustering) based on post-percentile normalized and mRNA noise removed datasets; and (**c**) SC3 cluster integration into PCA and *t*-SNE (upper panel) and gene marker discovery (lower panel). Silhouette plot, a quantitative measure of the diagonality of the SC3 consensus matrix, is based on *k*-means clustering. Known regulatory genes of pluripotent states, integrable with SC3 consensus clusters, are used as controls to validate the above meta-analysis approach and to differentiate stem cell states.

### Integration of RNA-seq data with microarrays for meta-analysis

To further validate the percentile normalization method, we used corroborated intra-laboratory datasets (D22B, D23, and D24) from Dr. Austin Smith’s laboratory, which were related to RNA-seq and microarrays (Supplementary Tables 1 and 2). These closely related datasets serve as reasonable controls to validate our normalization methods.

We initially calculated the correlation coefficient (*R*) between D22B and D23 or D24 cell samples pre- and post-percentile normalization with the inclusion of all possible sample pairs for comparison (Supplementary Fig. 1). We found that the correlation (R_pre_ = 0.55) between D22B (H9 cell line, RNA-seq) and D23 (H9 cell line, microarray) gene expression was significantly increased in post-percentile normalized datasets (*R*_post_ = 0.80, *R*_post_ - *R*_pre_ = 0.25; Welch modified *t-*test, *P*-value = 7.2E-37). Similar results were also found when comparing H9 Reset cell lines (Supplementary Figure 1). Evidently, there was a high pre-percentile correlation (*R*_pre_ = 0.87) between D22B (H9 Reset cell lines, RNA-seq) and D24 (HNES1-3, RNA-seq datasets). However, this correlation was actually decreased (*R*_post_ = 0.79, *R*_post_ - *R*_pre_ = −0.082; Welch modified *t*-test, *P*-value = 4.0E-16) post-percentile normalization between these RNA-seq datasets (Supplementary Fig. 1). Thus, these results suggest that some RNA-seq processing methods may overdrive the correlation of these datasets, which can also be attenuated by our percentile normalization.

Furthermore, between all naive-like cell lines, the mean correlation of pre-percentile normalized datasets (*R*_pre_ = 0.73, n = 7192 pairs,) was also significantly increased post-percentile normalization (*R*_post_ = 0.78; Welch modified *t*-test, *P*-value = 4.5E-50). Interestingly, between all primed cell lines, the mean correlation of pre-percentile normalized datasets was high (*R*_pre_ = 0.76, n = 35,370 pairs), which was slightly elevated post-percentile normalization (*R*_post_ = 0.79; Welch modified *t*-test, *P*-value = 3.4E-93). Thus, there was a similar correlation improvement for both naive-like and primed cell lines, suggesting that our analytic methods were appropriate for the assessment of pluripotent states in this study.

To further verify the comparability between RNA-seq and microarray gene expression, we performed PCA based on Pearson correlation (i.e., correlation-PCA or PCA hereafter) and focused on human oocytes and early embryos (2-cell to 8-cell stages and morulae), which require no (for oocytes) or minimal cell culture (for early embryos) and negligible laboratory protocol differences (Supplementary Table 2). PCA revealed that these oocytes and human early embryos were grouped or overlapped in the three major principal components (e.g., PC1, PC2, and PC3), particularly in PC2 that depicts nearly no variability in gene expression (Fig. 2, a1-3; b1-3). Taken together, the post-percentile normalization method followed by the exclusion of mRNA noise significantly improves the comparability between RNA-seq and microarray datasets, which may be suitable for analysis of datasets from different platforms in one meta-analysis.

**Figure 2.**
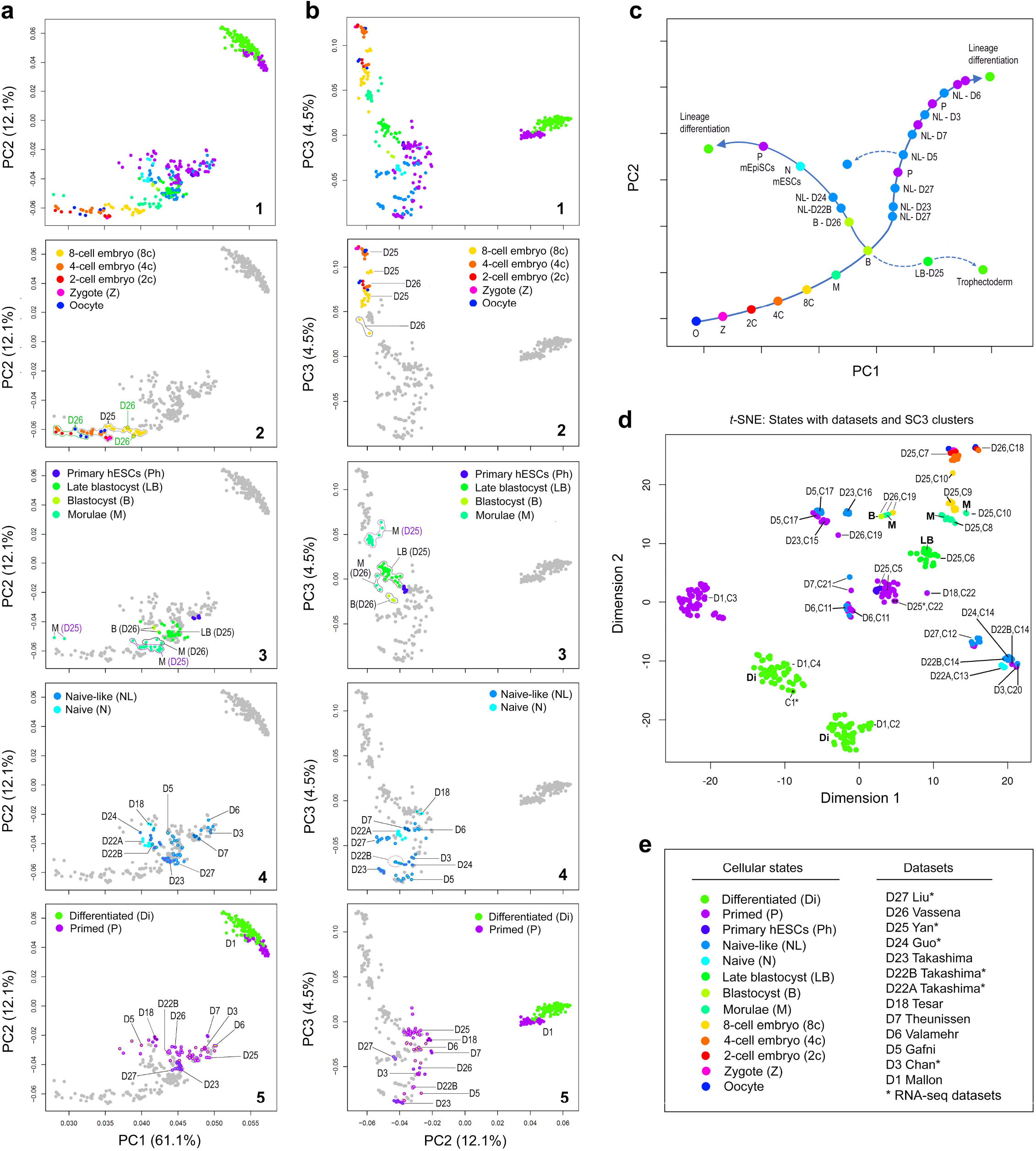
Meta-analysis combining PCA, *t*-SNE, and SC3 consensus clustering for accurately defining pluripotent and cellular states. The 13 datasets, used in this study, were named based on the first authors of the published reports. The datasets are composed of 434 samples from 10 independent laboratories, which can be identified with GSE and EMBL-EBI, and Sequence Read Archive (SRA) accession numbers in parentheses: D1 (GSE32923), D3 (E-MTAB-2031), D5 (GSE46872), D6 (GSE50868), D7 (GSE59435), D18 (GSE7866), D22A (E-MTAB-2857), D22B (E-MTAB-2857), D23 (E-MTAB-2856), D24 (E-MTAB-4461), D25 (GSE36552), and D26 (GSE29397), and D27 (SRP115256). (**a, b**) Correlation PCA. (**c**) The diagram illustrates the dataset-cellular state relationship observed (with solid curves) in both PC1 and PC2. Late blastocytes in D25 (LB-D25), indicated by a broken curve, contain heterogeneous cell types, in which some cells have a hypothetical trophectodermal fate. (**d**) A *t*-SNE visualization plot of all datasets with SC3 clusters and dataset numbers labeled. (**e**) Cellular states (with color keys) and datasets. Abbreviations: 2c, 4c, and 8c: 2-cell, 4-cell, 8-cell human embryos, respectively; B, blastocyst; D, dataset (e.g., D1, dataset 1); Di, differentiated hESCs; LB, late blastocysts; M, morulae; N, naive state; NL, naive-like state(s); O, oocyte; P, primed state; Ph, primary hESCs; Z, zygote.

### Unsupervised PCA, *t*-SNE, and SC3 clustering revealed interlaboratory data variations

By integrating genome-wide microarray and RNA-seq data into PCA, *t*-SNE, and SC3 clustering, we examined the relationship among all datasets, cell types, and cellular states in 2D plots. One of the most intriguing findings is that the interlaboratory data variations in some datasets, revealed by different principal components (e.g., PC1, PC2, and PC3) (Fig. 2a, 2b), are much higher than those of intra-laboratory data with respect to cell types, pluripotent states, and gene marker clusters. Generally, the majority of datasets within in a laboratory tend to group closer (e.g., D1, D3, and D6), regardless of the differences in their cell types. It is important to note these interlaboratory differences are likely due to adapted biological changes from cell culture protocols and practice peculiar to individual laboratories (Supplementary Tables 1 and 2), which may influence the interpretation of data concerning the pluripotent states.

Nonetheless, with our rigorous normalization approach, we were able to reveal a progressive cellular state transition from 2- and 4-cell human embryos toward 8-cell embryos, morulae, blastocysts, naive-like, and primed pluripotent states (Fig. 2a-c). Interestingly, there are two naive-like states, which are apparently associated with either naive mESCs or early blastocysts (Fig. 2a-c). For example, naive-like cells in D27, closely associated with those in D23, are adjacent to human blastocytes. Whereas the naive-like cells in both D22 B and D24 are grouped with naive mESCs (D22A and D18) (Fig. 2a4, b4, c). Moreover, the naive-like cells in D7, D3, and D6 are sequentially apart from both D27 and D23 in both PC1 and PC2, indistinguishing from some of primed hPSCs (Fig. 2a). Thus, these data indicate that naive-like and primed states can be definable in some datasets via meta-analysis. There are likely two distinct naive-like states based on their similarities to either mESCs or human early blastocysts. Collectively, our meta-analysis suggests a significant difference between interlaboratory naive-like and primed states of hPSCs.

To better visualize the results, we also performed *t*-SNE for the same datasets. Noteworthy, the D1 dataset, which represents one primed and two differentiation states (Supplementary Table 1), cannot be separated from one major group in Figure 2a. It can be visualized now as three distinct groups as expected in the *t*-SNE plot (Fig. 2d). Again, data or datasets within in the same laboratory incline to group closer in the *t*-SNE plot, regardless of the differences in their cellular states (Fig. 2d). To provide a better resolution of cellular or pluripotent states, we also employed transcriptomic cluster analysis (i.e., SC3 consensus clustering) based on the expression of all unique genes. SC3 clustering generated 22 consensus clusters (i.e., C1 to C22), in which 21 clusters enable the assignment of gene marker similarities to different pluripotent states of the cells (Fig. 2d). Noticeably, a single cluster (e.g., C7) defines multiple cellular states (e.g., oocytes, zygote, 2-cell and 4-cell embryos) in D25 (Fig. 2d). Conversely, multiple clusters (e.g., C5 and C22) may also define a specific pluripotent state (e.g., primed in D25) (Fig. 2c). Therefore, SC3 clustering combined with *t*-SNE and PCA likely offer a precise way to assess cellular states for large interlaboratory datasets, which have significant systemic variations.

### SC3 clustering unveiled multiple clusters that define various cellular and pluripotent states

Based on the 13 datasets, we have constructed a heatmap that is composed of the consensus clusters described above (Fig. 2d, Supplementary Fig. 2). We identified 22 individual gene clusters among these datasets after post-percentile normalization (followed by the mRNA noise exclusion method) using the SC3 clustering method (Kiselev et al., 2017). The dendrogram delineates the similarities or dissimilarities among 22 SC3 gene clusters, which are defined by a *P* value (*P* < 0.05) and the area under receiver operating characteristic (AUROC) (AUROC > 0.80) (Fig. 3a, 3b, Supplementary Table 7). The numbers of gene markers in all clusters range from 0 to 1252, with total 5,573 gene markers among the 22 clusters (Fig. 3b, Supplementary Table 7).

**Figure 3.**
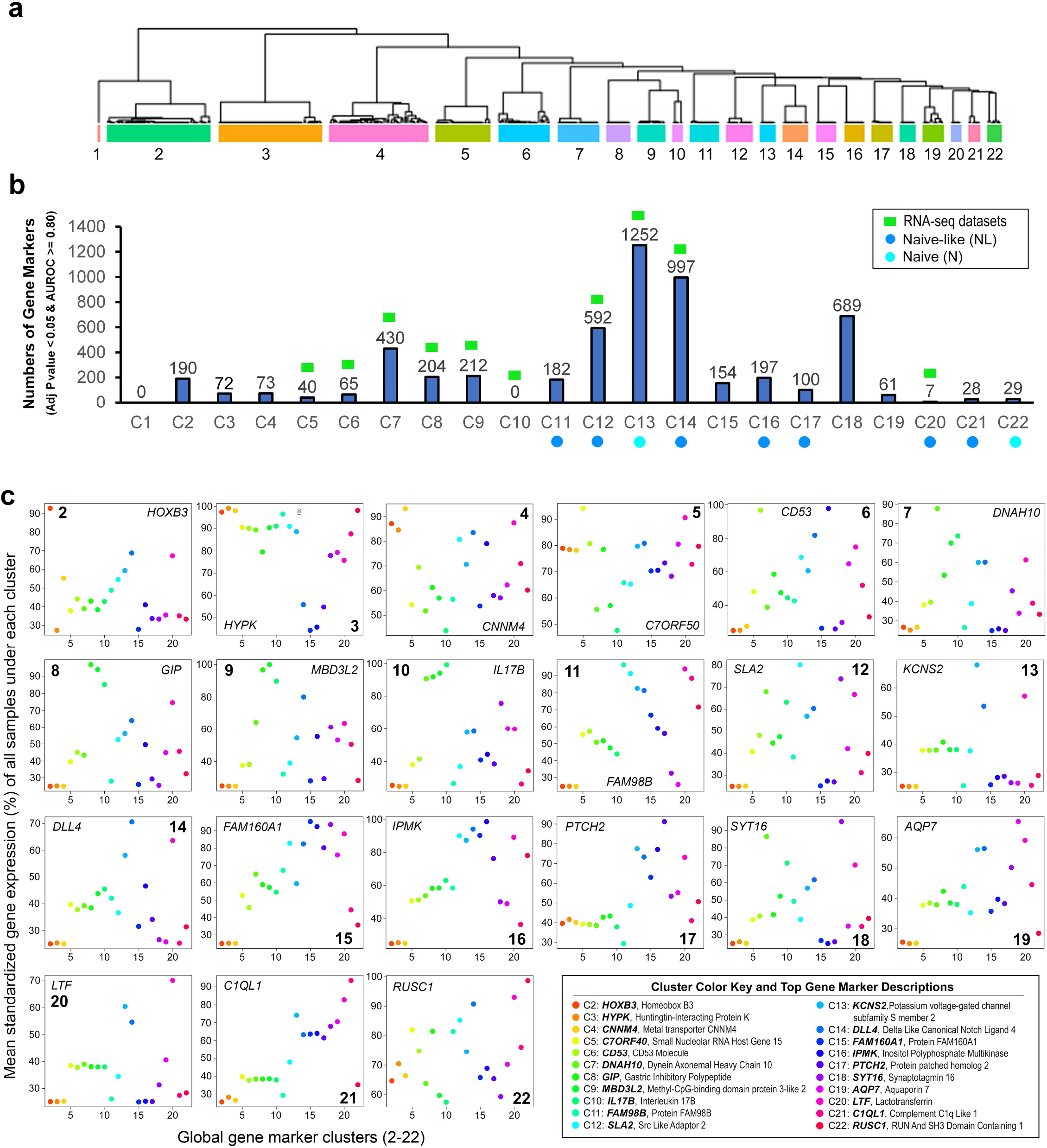
Transcriptomic clustering, gene marker identifications, and intercellular relationships among various independent studies. (**a**) Twenty-two SC3 clusters, derived from the post-percentile ranked and mRNA noise excluded datasets, were based on Euclidean, Pearson, and Spearman using SC3 consensus clustering. Shown here is the dendrogram delineating the similarities among 22 SC3 gene expression clusters. The whole heatmap is available in Supplementary Figure 2. (**b**) Histogram that summarizes the numbers of gene markers in 22 SC3 clusters, which are defined by *P* values (< 0.05) and the area under receiver operating characteristic (AUROC > 0.80). Light blue- and cyan-colored dots are used to denote the clusters defining naive-like and naive states, respectively. For better visualization, RNA-seq information is labeled on the top of the box plot. Abbreviations: Adj, adjusted. (**c**) Top gene marker presentation in the 21 unsupervised individual clusters. Each colored dot represents mean standardized gene expression (>25^th^ percentile) of all samples per cluster. Only the top 1 gene marker is labeled in the plot.

To evaluate the relative specificity of top gene markers among the 21 clusters, we calculated the mean standardized expression (ranging 1-100%) of each gene marker in all samples under each cluster. It appears that the top gene marker expression displays a significant fluctuation/scatter among the 21 clusters (Fig. 3c). It is yet unknown how the vast majority of these top gene markers in each individual cluster regulate the fates of pluripotent stem cells both *in vivo* and *in vitro*. Hence, we unbiasedly surveyed the top 15 gene markers in each individual cluster of interest (Supplementary Table 7), with a focus on those having curated known functions in reliable informatics (www.ncbi.nlm.nih.gov, www.genecards.org, www.uniprot.org). Collectively, there are 7 gene clusters (i.e., C11, C12, C14, C16, C17, C20, and C21), characteristic of *in vitro* cell types with naive-like states, and 2 clusters (C13 and C22) associated with naive states (Fig. 3b, Supplementary Table 7).

With regard to SC3 clusters defining the datasets with naive-like states, C11, characteristic of both primed and naive-like cells in D6, possesses several gene markers of interest, which are involved in the mTORC1 pathway (e.g., *RHEB*), the formation of the mitochondrial complex III (e.g., *TTC19*), binding of membrane phospholipids (e.g., *FGD2*), and the maintenance of the epithelial feature (e.g., *ITGAE as a* coreceptor gene for E-cadherin). C12 defines both primed and naive-like cells in D27. There are at least several genes known for their roles in transcriptional regulation (e.g., *LIN54*), histone-binding and nucleosome-remodeling (e.g., *BPTF* and *MLLT10*), and cell-cycle checkpoint control (e.g., *DONSON*). The *LEO* gene is of interest. It encodes the RNA polymerase-associated protein LEO1, which is a component of the multiple functional PAF1 complex (PAF1C) implicated in the regulation of embryonic stem cell pluripotency.

C14 delineates both naive-like and primed states in D22B. The functional roles of the top 15 gene markers (e.g., *ZFP30*, *ANKRD34A*, and *FAM78B*) in the regulation of pluripotency and differentiation are unknown. However, at least, 40% (6/15) of the top gene markers (e.g., *ASCL1, CNTF, DLL4, KCNA6, WNT9A,* and *ZBTB18*) are associated with neuronal functions and brain development, suggesting a propensity for neuroectodermal differentiation. C16 characterizes a gene cluster specific to the naive-like state in D23. The top gene markers in this cluster embrace *IPMK* (inositol polyphosphate multikinase required for normal embryonic development) and *HRK* (encoding Harakiri, a BCL2 interacting protein that promotes apoptosis). At least 20% (3/15) gene markers are implicated in mitochondrial functions, including *TIMMDC1* (involved in the assembly of the mitochondrial Complex I), *CKMT1A*/*CKMT1B* (mitochondrial creatine kinase genes), and *FXN* (that plays a role in mitochondrial iron transport and respiration).

C17 specifies both naive-like and primed states in D5. Clearly, 2 gene markers (e.g., *AKT1* and *PTCH2*) are involved in well elucidated signaling transduction pathways. More than 25% of the gene markers are associated with diverse metabolic processes, including ferrous iron binding and oxidative RNA demethylase activity (mediated by *FTO*), the conversion of N-acetyl-D-glucosamine to N-acetyl-D-glucosamine 6-phosphate (by *NAGK*), the transfer of sulfate from 3’-phosphoadenosine 5’-phosphosulfate to nitrogen of glucosamine in heparan sulfate (by *NDST1*), and protein and sphingolipid metabolism (by *SUMF2*).

C20 only has 7 gene markers that are associated with both naive-like and primed states in D3. Measurably, 2 gene markers are shown to participate in the regulation of cellular growth and differentiation (e.g. *LTF*) and controlling homeostasis (e.g., *ADCY8*). Approximately, 29% of them (e.g., *FAM83C* and *INSRR*) function in the MAPK-ERK signaling pathways that directly control mammalian ESC pluripotency. Lastly, C21 elucidates both naive-like and primed states in D3. Nearly, 40% (6/15) of the top gene markers (i.e., *NDUFB3, EPN1, PI4KA, NDUFA11, NDUFC1,* and *DGKZ*) are associated with metabolism. Particularly, 20% of them (i.e., *NDUFB3*, *NDUFA11*, and *NDUFC1*) are parts of mitochondrial NADH dehydrogenase (Complex I).

With regard to SC3 clusters defining the datasets with naive states, C13 represents an independent cluster that define naive mESCs in D22A. Interestingly, this cluster is highlighted by the zinc finger E-box binding homeobox 1 (*ZEB1*) with two metabolism-related gene markers (*PADI4* and *HK3*). Importantly, a newly identified naive regulatory gene *GJB5* in mESCs, which encodes the gap junction beta-5 protein, is also within the top 15 gene cluster. Furthermore, C22 defines both naive mESCs and primed mouse EpiSCs in D18 as well as 2 primed hESCs from D25. A cluster of energy metabolism genes is highlighted: *COX17* (Cytochrome C oxidase copper chaperone *COX17*), *ETFB* (electron transfer flavoprotein subunit beta), and *NDUFA7* (NADH:ubiquinone oxidoreductase subunit A7).

In summary, SC3 clustering readily identifies potential regulators and mediators (e.g., FAM83C, GJB5, INSRR, IPMK, and LEO), which control embryonic development and embryonic stem cell pluripotency. Expression of a notable cluster of genes (e.g., *CKMT1A/CKMT1B, COX17, ETFB, FXN*, *NDUFA7, NDUFA11*, *NDUFB3*, *NDUFC1, TIMMDC1*, and *TTC19*) highlights the demand of mitochondrial activities including oxidative phosphorylation in these hPSCs. Moreover, expression of top gene markers in individual clusters is frequently associated with the regulation of cell death and survival (e.g., *RHEB, HRK, AKT1,* and *PTCH2*), cell-cycle checkpoint control (e.g., *DONSON*), generic metabolic processes (e.g., *DGKZ, NAGK*, *NDST1*, *PI4KA*, and *SUMF2*), and regulation of gene expression (e.g., *BPTF, LIN54,* and *MLLT10*). Thus, the relative specificity of top gene marker expression would enable its utility to define cellular identities, states, and potential biological functions in NLPs that are generated from different laboratories.

### Tabular tools for predicting cellular states

To reveal the complicated relationship between different cell types (e.g., oocyte and zygote), human early embryos (2-cell to late blastocysts), pluripotent (e.g., naive, naive-like, and primed), and differentiated states, we linked cellular state data to both SC3 clusters and the cellular identities from the 13 datasets. As shown in Figure 4, all cellular and pluripotent states can be assigned into all 22 clusters. Furthermore, this comparative analysis confirms the significant heterogeneity among naive, naive-like, and primed states (Fig. 4). For examples, the naive mESCs from both D18 and D22A clustered significantly differently. There were two naive-like states. One was associated with the naive mESCs in D22A, which were cultured under hypoxia and processed by the RNA-seq platform. Another one was linked to the naive mESCs in D18, which were cultured under normoxia and processed by cDNA microarray (Fig. 4). Thus, the cluster-dataset relationship in these specific sample classes may be explained by the use of different culture methods. However, we cannot rule out that the laboratory protocols used for analyzing RNA expression might also contribute to data variability.

**Figure 4.**
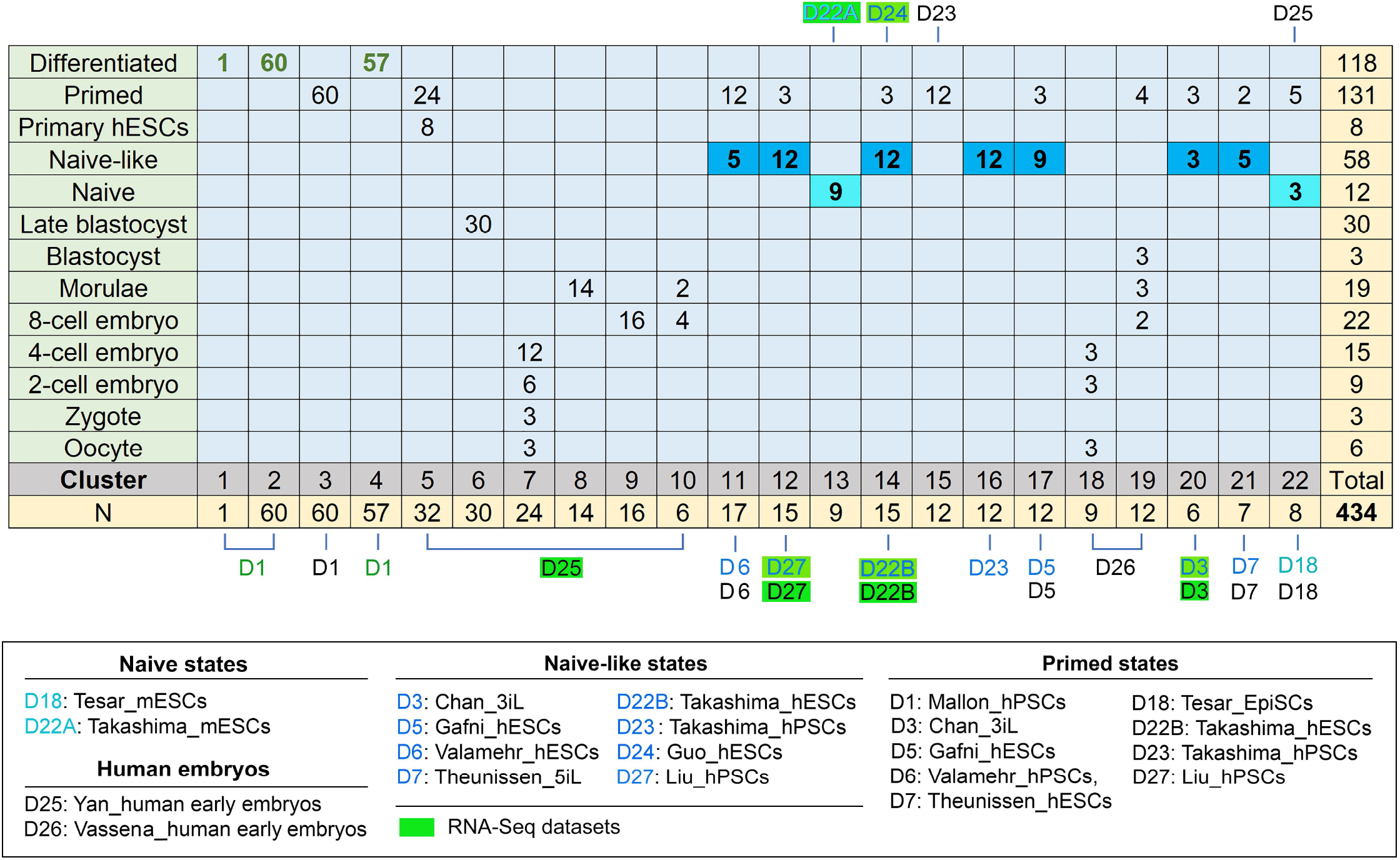
Cluster-state-dataset associations. Tabular tools for predicting the relationships between different cellular (e.g., oocyte and zygote), pluripotent (e.g., naïve, naïve-like, and primed), and differentiated states, all of which are associated with the 22 SC3 clusters in 13 datasets. To interpret the expression of different gene clusters: if we look at differential expression between cell states (e.g., naive-like and primed) in one dataset (e.g. D6) and compare to differential expression (i.e., naive-like and primed) in a second dataset (e.g., D5). The result appears to be quite different: D6 and D5 are separated by 6 SC3 clusters, which might suggest a major difference in their actual cellular states. Lower panel: detailed descriptions of pluripotent states that are associated with individual datasets, in which mESCs are used as naive state controls for facilitating comparative analysis. Of note, one dataset may contain two or more different states, which are distinguished by a different font color. For example, 22B is labeled with two different colors (black and blue), in which 22B in light blue font denotes the naive-like state, whereas 22B in back designates the primed state. Abbreviations: N, the number of samples used in each cluster; Naive, naive pluripotent state; Naive-like, naive-like pluripotent state; Primed, primed pluripotent state.

Moreover, naive and naive-like state clusters do not overlap, suggesting that hPSCs with naive and naive-like states have different pluripotent states. Within the datasets D3, D5, D6, D7, D22B, and D27, the cell samples with naive-like states share clusters with primed states, suggesting that these naive-like hPSCs have a closer relationship with primed hPSCs than naive mESCs. In contrast, naive-like and primed states in D23 were clustered separately by C15 and C16 respectively (Fig. 4), suggesting that the naive-like state in D23 is indeed significantly different from the primed state within the same dataset. This results also demonstrate that our analytic approach can differentiate between naive-like and primed states. Noticeably, the above distinctions have not been revealed previously, which might be useful for defining accurate pluripotent states in hPSCs.

### Supervised cluster analysis defining new gene signatures for characterizing pluripotent and cellular states

Beside the above unsupervised meta-analysis, we also present here 4 supervised heatmaps that delineate normalized mean expression of 68 known genes (www.genecards.org), which encode dominant signaling molecules (n = 23, Fig. 5a), differentiation markers (n = 17, Fig. 5b), developmental regulators (n = 18, Fig. 5c), and key pluripotency transcriptional factors (n = 10, Fig. 4d). These heatmaps demonstrate new features of cellular and pluripotent similarities associated with the 21 SC3 gene clusters as described in Figure 3a.

**Figure 5.**
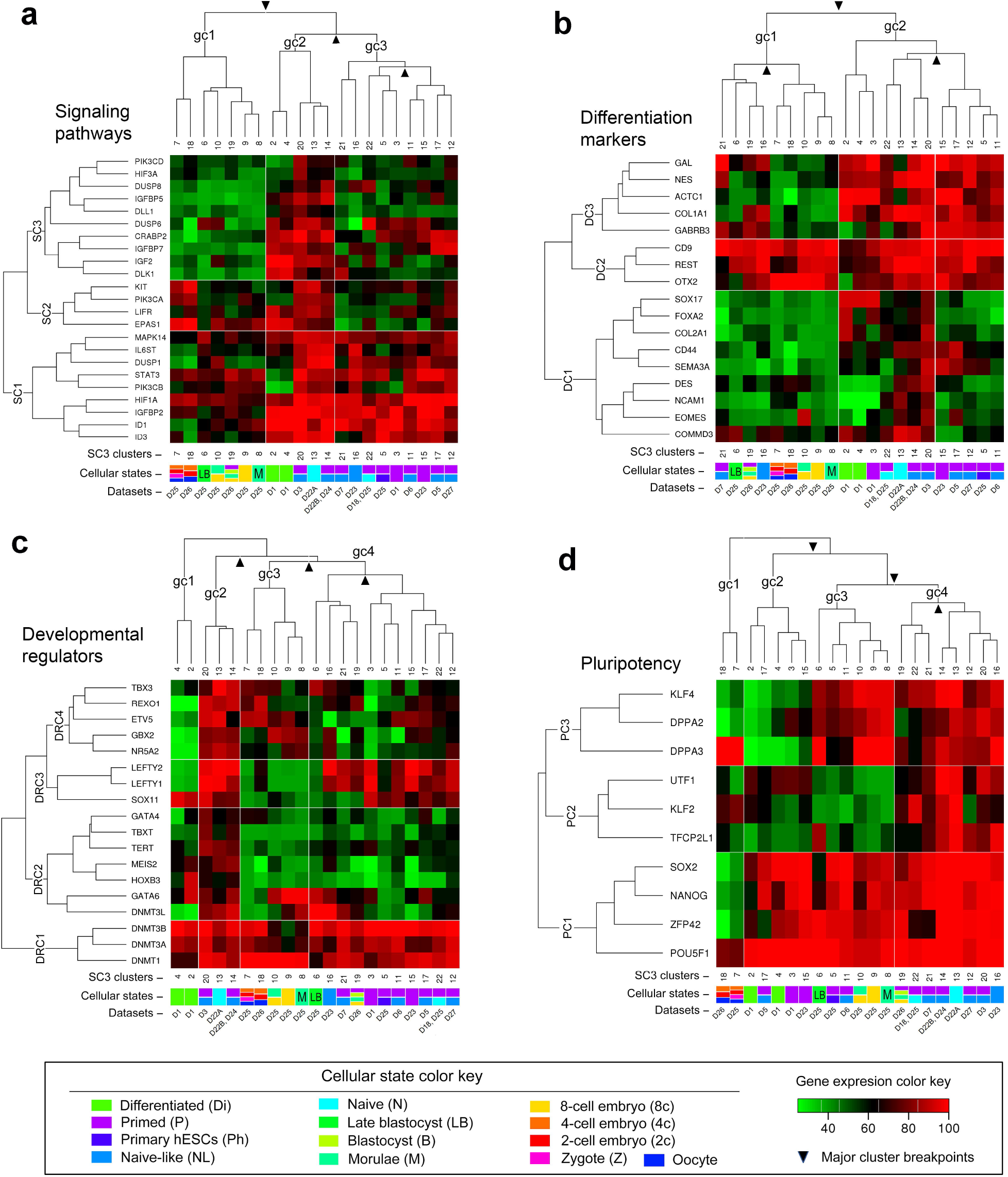
Supervised gene cluster analysis of cellular states. Heatmaps are generated with normalized mean expression for supervised gene markers from all samples under each defined cluster. Shown here are the heatmaps of normalized mean gene expression (>25^th^-100^th^ percentile) across all clusters for supervised (known) gene markers, in which dominant signaling pathways (n = 23), differentiation marker (n = 17), developmental regulators (n = 18), and pluripotency transcriptional factors (n = 10) were used for the process of SC3 clustering algorithms. Of note, unsupervised SC3 employs 5,573 unique gene markers for clustering analysis (Fig. 3a, b; Supplementary Table 7). Both cellular states and datasets, which are associated with SC3 clusters, are labeled on the bottom of each heatmap. Each individual cluster contains one or more cell types or states as indicated by color coding detailed in the lower panel. The number of gene markers used for conducting unsupervised or supervised clustering analysis has a significant impact on the relationship among various cellular and pluripotent states as shown in these heatmaps. For details, see Supplementary Table 7. Additional abbreviations: DC, differentiation gene cluster; DRC, developmental regulator gene cluster; gc, global cell cluster; PC, Pluripotency gene cluster; SC, signaling pathway cluster.

We initially analyzed the influence of gene markers that encode common signaling pathways. As indicated by annotated major cluster breakpoints, mean standardized expression of these markers results in three major signaling gene clusters (SC1, SC2, and SC3) that classify the 21 clusters into three global cell clusters (gc1, gc2, and gc3) (Fig. 5a). The gc1 contains 7 SC3 clusters, defining human oocytes and early embryos by Vassena and Yan datasets (Vassena et al., 2011; Yan et al., 2013). Interestingly, upregulation of the 4-gene subset (i.e., *KIT, PIK3CA, LIFR,* and *EPAS1*) is a strong indicator for the mixed cellular states that contain oocytes, 2-cell and 4-cell embryos. Clearly, all naive, naive-like, and primed clusters appear to be driven by the expression of a 9-gene cluster (*MAPK14, IL6ST, DUSP1, STAT3, PIK3CB, HIF1A, IGFBP2, ID1,* and *ID3*) in the SC1 block (Fig. 5a). The naive and primed datasets (D3, D22B, and D24) share the 23-gene cluster with the naive mESCs in D22A (Fig. 5a), suggesting the closest relationship among the three datasets from two different laboratories when only considering this signaling transcriptomic signature. Nonetheless, these signaling gene markers do not provide clear signatures to distinguish naive from naive-like and primed states.

We next examined the impact of differentiation gene markers on the rearrangements of the 21 clusters. As shown by the cluster breakpoints, mean standardized expression of these gene markers results in three major differentiation clusters (DC1, DC2, and DC3) that organize the 21 SC3 clusters into two global cell clusters (i.e., gc1 and gc2) (Fig. 5b). Under this analysis, DC1 can cluster all naive, naive-like, and primed states into gc2 (Fig. 5b). Moreover, the two clusters (C13 and C22), containing the two mESC datasets, bear most resemblance to each other under this supervised condition (Fig. 5b), which contrasts with our unsupervised analyses.

Concerning the influence of developmental regulators on the relocations of SC3 clusters, three major DNA methyltransferase genes (i.e., *DNMT1, DNMT3A,* and *DNMT3B*) showed ubiquitously high levels of mRNA expression in all clusters (Fig. 5c), thus diminishing their predictive values for the pluripotent states. However, the predictive value for both naive and naive-like states might underlie the expression patterns of *TBX3, REXO1, ETV5, GBX2, and NR5A2* within the DRC4 cluster and of *DNMT3L* and *GATA6* within DRC2 (Fig. 5c). Again, the three naive and primed datasets (D3, D22B, and D24) share the entire 23-gene cluster with the naive mESCs in D22A (Fig. 5c), similar to the finding observed in signaling pathways (Fig. 5a). Of note, the *Xist* transcripts, one of the developmental hallmarks of naive pluripotency, was not included in this analysis due to the mRNA noise exclusion step.

Finally, we interrogated the effect of known pluripotent regulators on gene cluster rearrangements. We found that three well-established pluripotent regulator genes (i.e., *SOX2, NANOG, and POUF51*), clustered with a poorly understood regulatory gene *ZFP42* (PC1), had no predictive values for discriminating diverse pluripotent states (Fig. 5d). However, the predictive values for both naive and naive-like states clearly underlie the expression pattern of the PC2 (*UTF1, KLF2,* and *TFCP2L1,*) and PC3 (*KLF4, DPPA2,* and *DPPA3*) clusters (Fig. 5d). Moreover, the 3-gene cluster *UTF1, KLF2,* and *TFCP2L1* discriminates mESCs (C13 and C22) from human early embryos and other naive-like states. Thus, these supervised analyses not only confirm the value of some previously identified gene markers (such as *KLF2* and *TFCP2L1*) but also validate our normalization approach to discern pluripotent states.

## DISCUSSION

Achieving human naive pluripotency through the perturbation of growth factor signaling is believed to have significant impacts on hPSC growth, expansion, genetic engineering, disease modeling, and drug discovery. However, to accurately define a pluripotent state seems to be hindered by a lack of reliable analytic tools for comparative meta-analysis. Based on the integrated meta-analysis described in this study, we have provided an unbiased assessment of cellular and pluripotent states. Accordingly, we have integrated PCA and *t*-SNE with 21 SC3 clusters that define various cellular states, including enriched panels of regulatory, metabolic, and effector gene markers (Figs. 3, 4, Supplementary Table 7). Here, we will discuss critical interference factors of multivariate meta-analysis, the rationale and reliability of our analytic approach, integration of supervised into unsupervised clustering analyses, and naive hPSC growth properties under certain defined growth conditions.

The purpose of meta-analysis is to mitigate the interference of laboratory-specific batch effects while preserving genuine biological differences across all datasets. Batch effects, unwanted variations attributable to technical sources, are common in high-throughput biology, which substantially confound meta-analysis across different datasets. Hence, beside the interlaboratory protocol differences (Supplementary Table 2) that may derail a successful meta-analysis, other influencing batch factors (e.g. cDNA microarray and RNA-seq) should also be taken into consideration in this study. For example, when dealing with cDNA microarray and RNA-seq data for the analysis, we frequently encounter: (i) sensitivity related to Poly(A) in channel versus ribosomal RNA depletion; (ii) sequencing depth (e.g., coverage, numbers of reads, and the length of reads used for mapping); (ii) paired versus single ends (in which paired ends are more accurate); (iv) stranded versus non-stranded; (v) outliers, and importantly, (vi) low levels of mRNA noise. Accordingly, the distinct SC3 clustering differences (C6-10 versus C18-19) between the human early embryo datasets D25 (RNA-seq) and D26 (cDNA microarray) seem to be consequential to one of the above discussed issues (Fig. 4).

Both ComBat and SVA (surrogate variable analysis) are the popular methods used to correct the batch effects due to their high performance across different platforms (Jaffe et al., 2015; Johnson et al., 2007; Leek et al., 2012). However, these batch correction methods may not be suitable for our datasets by design due to the existence of numerous limitations (Buhule et al., 2014; Goh et al., 2017; Jaffe et al., 2015; Kupfer et al., 2012; Leek and Storey, 2007; Nygaard et al., 2016). In the case of this study, our collected datasets represent unique sample types, passages, protocols, and are therefore mutually exclusive. Neither are batch sizes equal nor are sample classes evenly distributed across datasets (batch groups). Under this unbalanced situation, applying batch-effect correction may increase false positive discoveries (Nygaard et al., 2016). Specifically, SVA may reduce intra-class variability at the cost of losing sample-specific information (e.g., subpopulation effects) (Jaffe et al., 2015). Subpopulations, typically under-represented in datasets, are difficult to identify, which may be of biologically valuable for *in vitro* cell culture. Subpopulation effects can be removed due to that they are similar to batch effects (Leek and Storey, 2007).

The rationale of using meta-analysis relies on validated normalization strategies that integrate interlaboratory datasets generated from different platforms. To enable impartial comparison of the transcriptomic levels of interlaboratory datasets, we demonstrate one such pre-processing strategy that involves quantile normalization of mRNA expression across samples within each dataset, percentile coding, and subsequent mRNA noise exclusion. Thus, our approach does not batch correct the data. Rather, we use percentile coding to preserve relative expression abundance (ranking) within the sample. It is these rankings that can then be directly and fairly compared across datasets. Our analytic approach is similar to that Gene Fuzzy Scoring (GFS), an emerging powerful data normalization method against batch effects (Belorkar and Wong, 2016). By GFS, gene features are ranked and assigned with new values (ranging from 0 to 1) based on high, moderate, and low confidence gene features. The low confidence features are considered as noise and penalized by flooring the expression values to zero, which is similar to the analytic methods that we described in this study. Recently, Wells and colleagues also used similar percentile normalization but without the exclusion of mRNA noise step. They successfully facilitated the comparison of different experimental series of myeloid cells in the Stemformatics platform (doi.org/10.1101/719237). Thus, our method enables us to eliminate large amounts of unwanted variations and to boost high confidence signal. With our rigorous approach, the differences between naive-like, primed, and differentiated states remain preserved (Fig. 2–5). Hence, the relationship between various cellular states is clearly definable (Fig. 6). Moreover, the transformed values for each mRNA can be reanalyzed and representative markers for each cluster identified, including those that discriminate the pluripotent state (e.g., primed and naive).

**Figure 6.**
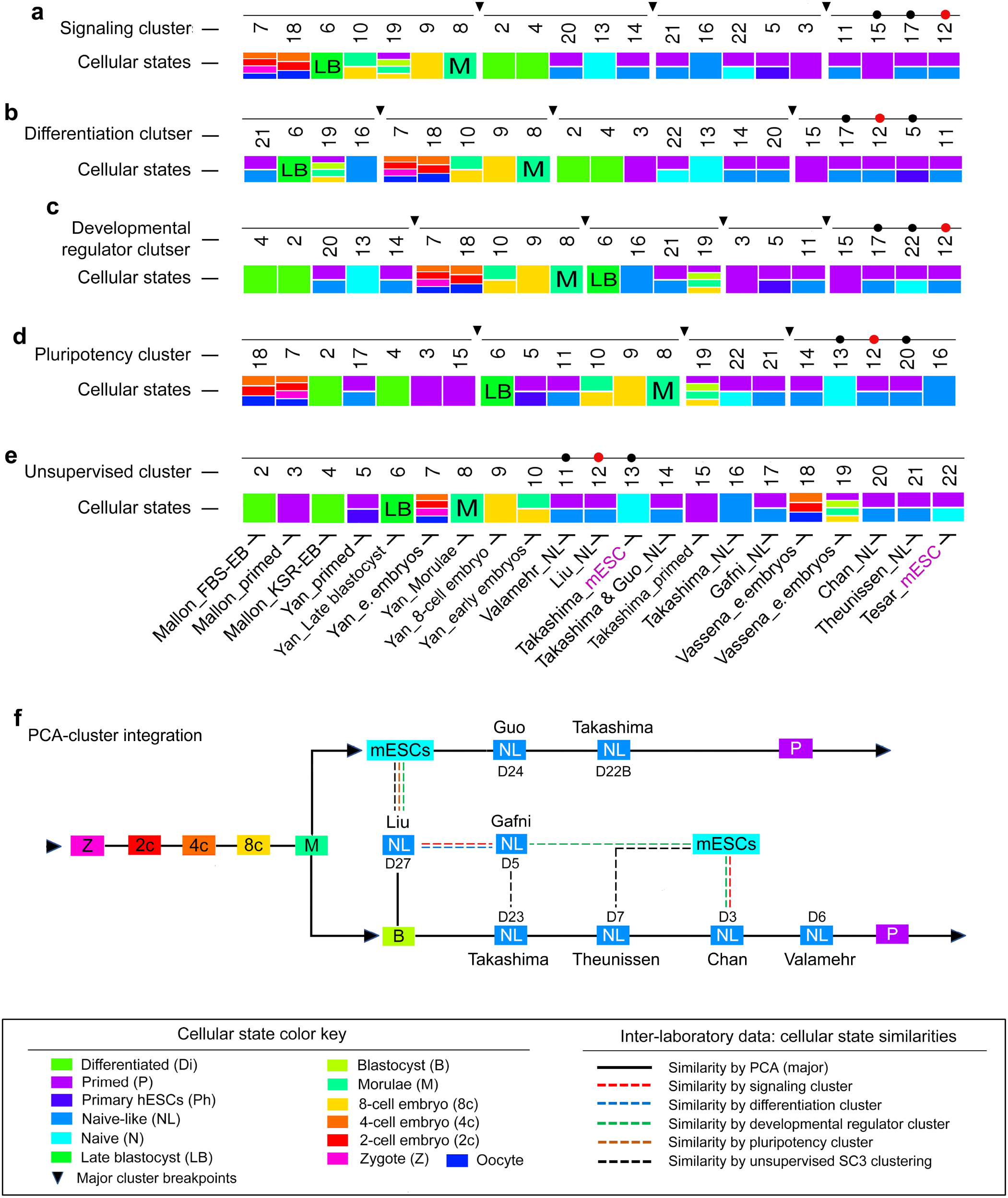
Integrated analysis of cellular states. (**a-e**) One-dimensional mapping of pluripotent and cellular identities by integrating cellular states with supervised (n = 4) and unsupervised SC3 cluster analysis. Each individual cluster contains one or more cell types or states as indicated by color coding detailed in the lower panel. The clusters and cellular states are linked to the datasets that contain human early embryos, naive mESCs, naive-like hPSCs, and differentiated EBs from hPSCs. Black arrowheads indicate cluster breakpoints that enable cluster rearrangements for better visualization of the cluster relationship. The three dots labeled on each cluster are used to trace the relationship between the clusters under different clustering conditions. (**f**) Integration of PCA into SC3 clustering analysis. The diagram delineates the similarities between naive-like states from different laboratories. Human blastocysts (B) and naive mESCs are used markers to depict the above relationship. The relationship is mainly based on PCA (as connected by the black solid lines) and partially on SC3 clusters (connected by one or more broken color lines as detailed in lower panel). Additional abbreviations: e., early; FBS_EB, fetal bovine serum (FBS)-mediated embryoid body (EB) differentiation; KSR_EB, KnockOut Serum Replacement (KSR)-mediated embryoid body (EB) differentiation; mESC, naive mouse embryonic stem cells; NL, naive-like pluripotent state; Primed, primed pluripotent state.

The reliability of this analysis underlies its capacities to significantly increase the comparability between RNA-seq and microarray datasets without a substantial bias (Fig. 2–5, Supplementary Fig. 1) and to generate compelling gene markers that are highly relevant to pluripotent stem cell biology. Indeed, with our analytic approach, many meaningful new pluripotent markers or regulators have been identified in this analysis (Fig. 3 and 4; Supplementary Table 7). Some of these newly identified top gene markers enable us to understand how the diverse transcriptomic signatures orchestrate to achieve the complicated pluripotency dynamics. For example, *MLLT10*, identified from C12, encodes Protein AF-10 that binds to unmodified histone H3 and regulates DOT1L functions including histone H3K79 dimethylation (Chen et al., 2015). Evidently, MLLT10 physically interacts with the Oct-4 protein, a fundamental regulator of pluripotency, in purified Oct-4 protein complexes from naive mESCs (Ding et al., 2012), suggesting a potential role of the MLLT10-Oct-4 complex in the regulation of human naive pluripotency. Moreover, *GJB5*, identified from C13, encodes the Gap junction beta-5 protein that mediates cell-cell contacts and direct intracellular communication. Knockdown of *Gjb5* in mESCs results in downregulation of critical pluripotency genes, thereby leading to cellular differentiation (Bernardo et al., 2018). Thus, several gene markers identified in this analysis have already shed some light on the biochemical and epigenetic aspects of naive-like pluripotency. Moreover, top gene markers can be integrated into PCA to achieve multifaceted view of diverse pluripotent states.

Integration of supervised into unsupervised analyses would have *pros* and *cons* in this meta-analysis. The supervised approach may increase the sensitivity to reveal some well-known common features of hPSCs or NLPs (Fig. 5 and 6). However, it does not reflect cellular states at a genome-wide scale. Thus, a statement made from either a supervised (usually with a small subset of gene markers) or unsupervised analysis should be weighed differently. However, we may integrate supervised into unsupervised analyses in a one-dimensional format (Fig. 6a-e). This dimensional reduction provides a rapid, informative, and unbiased view of cellular and pluripotent states under various interlaboratory growth and assay conditions. For example, the naive mESCs from D18 showed the closest relationship (as indicated by C22) with the primed EpiSC control from the same dataset in clustering analysis. However, the two naive mESC controls (from D18 and D22A) do not show a closer relationship in PCA plots, *t*-SNE, and SC3 clustering (Fig. 2a-d, 3, and 4). This discrepancy between the two mESC lines may be explained by their different cell culture methods, in which the mESCs from D18 were cultivated under a normoxic condition (19% O_2_) whereas the mESCs from D22A were maintained in hypoxia (i.e., 5% O_2_).

Moreover, in another independent meta-analysis, Smith and colleagues showed human naive-like Reset cells (induced by 2iL and the PKC inhibitor Gö6893) were similar to their naive mESCs in PCA plots (Takashima et al., 2014). In their meta-analysis, they indicated that NLPs (from both Gafni and Chan datasets), located distal from the Takashima datasets in the PCA plot, are more similar to their primed counterparts (Takashima et al., 2014). Clearly, our meta-analysis revealed much larger differences between some interlaboratory datasets than their actual cellular state differences between intra-laboratory cell samples. Under these circumstances, the integration SC3 clusters into PCA may provide an unbiased view of the cellular state relationship between different laboratories (Fig. 6f). Our meta-analysis suggests the possibility that the previously reported similarities or differences in some naive and naive-like cellular models are likely attributed to interlaboratory protocol differences.

With respect to PCA, separation of interlaboratory data in the plots seems to be insufficient when including multi-laboratory datasets for the analysis. This problem may be overcome by comparing PCA plots with both *t*-SNE and SC3 clustering (Fig. 2–4). Concerning *t-*SNE visualization and cluster analysis, the high-dimensional data reduction technique, originally developed by van der Maaten and Hinton in 2008 (van der Maaten and Hinton, 2008), has gained popularity for data analysis and machine learning in recent years. The greatest advantage of *t*-SNE lies in its ability to visualize data in a fascinating 2D plot for high-dimensional datasets (up to thousands of dimensions). However, we should be aware that this technique is a random and non-linear method. Its dimensions, physical distances of data points, size of clusters are meaningless in the plots (https://distill.pub/2016/misread-tsne/). Noticeably, *t*-SNE plots may differ from each other in the same datasets. To enhance the strength of *t*-SNE in meta-analysis, we emphasize the combined use of *t-*SNE with both PCA and SC3 clustering as demonstrated in this study.

In summary, our meta-analysis indicates that hPSCs grown under current naive-like protocols do not have overlapping naive pluripotency clusters with those of mESCs. However, the naive-like hPSCs in some datasets (e.g., D23 and D27) show closer resemblance to either naive mESCs or human early blastocysts or both. Likely, there is considerable heterogeneity among various cellular and pluripotent states in a large cohort of datasets generated from different laboratories. Interlaboratory data variation represents the predominant factor interfering with the accuracy of meta-analysis, which has been under-evaluated in previous studies and thus significantly limits the predictive values for defining cellular and pluripotent states. The combined use of percentile normalization with PCA, *t*-SNE, and SC3 clustering, which represents a new strategy to compare multiple interlaboratory datasets, has significantly improved the predictive values of the current meta-analysis. Other data normalization or transformation algorithms, aiming to be resistant to the batch effect of interlaboratory samples, should also be considered in the future. It would be crucial for reducing interlaboratory data disparities.

## METHODS

### Datasets for meta-analysis

We collected 13 datasets for multivariate meta-analysis (Chan et al., 2013; Gafni et al., 2013; Guo et al., 2016; Liu et al., 2017; Mallon et al., 2013; Takashima et al., 2014; Tesar et al., 2007; Theunissen et al., 2016; Valamehr et al., 2014; Vassena et al., 2011; Yan et al., 2013). These datasets are composed of 434 samples from 10 independent laboratories (Supplementary Tables 1 and 2). The datasets can be identified with GSE and EMBL-EBI accession numbers in parentheses: D1 (GSE32923), D3 (E-MTAB-2031), D5 (GSE46872), D6 (GSE50868), D7 (GSE59435), D18 (GSE7866), D22A (E-MTAB-2857), D22B (E-MTAB-2857), D23 (E-MTAB-2856), D24 (E-MTAB-4461), D25 (GSE36552), D26 (GSE29397), and D27 (SRP115256). We curated these datasets based on their laboratories, first author(s), the size of samples (n), cell types (e.g., blastocysts and hESCs), cellular states (e.g., primed or naive-like), culture medium with growth factors, protocols, feeder/coating (e.g., MEFs or Matrigel), oxygen tension (e.g., normoxia and hypoxia), and RNA processing platforms (e.g., microarray and RNA-seq) (Supplementary Tables 1-3). Of note, there is a substantial difference in the cell culture protocols used to generate and maintain hPSCs between individual laboratories, particularly in the methods used to derive naive-like hPSC (Supplementary Tables 1-2). Another major protocol difference is the use of oxygen. For example, the naive and naive-like cells in datasets (D3, D6, and D18), were cultured under normoxia, which contradicts with the majority of these cells that were grown under hypoxia. Moreover, the primed hPSCs in three datasets (D3, D6, and D27) were maintained in either Matrigel (D3 and D6) or vitronectin (D27) coated plates. The remaining primed hPSCs and all naive or naive-like cells were cultured on MEF feeders. All detailed information can be found in Supplementary Tables 1-3.

### RNA-seq and microarray datasets for validation of normalization methods

We used validated intra-laboratory datasets from Dr. Austin Smith’s laboratory, which are related to cDNA microarray and RNA-seq. These datasets can be briefly described as follows: D22B (accession E-MTAB-2857, RNA-seq; primed H9 cell lines, n = 3; H9 Reset lines, n =3), D23 (accession E-MTAB-2856, microarray; primed H9 cell lines, n=3; H9 Reset lines, n = 9), and D24 (accession E-MTAB-4461, RNA-seq; HNES1-3 Reset cell lines, n = 9). More detailed information is available in Supplementary Tables 1-3.

### Data transformations for meta-analysis

Meta-analysis was used to analyze the above 13 datasets in order to compare the pluripotent and differentiation states. To enable different datasets to be used for meta-analysis, we transformed all datasets by the following major steps: (i) quantile normalization within datasets using the R code, (ii) data filtering to retain unique genes only, (iii) data collapse methods to calculate median values of multiple gene probes, and (iv) percentile coding (from 1 to 100%) of each gene expression in each sample followed by removing mutually expressed genes below the 25^th^ percentile (Supplementary Tables 5 and 6).

Briefly, we performed quantile normalization (Log_2_) of the transcriptome per dataset (total datasets = 13) and created a master expression matrix of 11,606 unique genes in rows and 434 samples in columns. The expression levels recorded in this matrix by all samples were further coded by quantile bins (1-100%). Explicitly, the observed expression values across all genes for a sample were used to define the 1^st^ to 100^th^ quantile values. These quantile values were then used to code where each expression value for the sample falls. For example, if an expression value for a gene of the sample falls between the 20^th^ and 30^th^ quantile values, the gene expression then has a defined value of 20. This quantile-bin approach was applied for all genes per individual sample.

By doing so, the ranking of RNA expression within a sample for a dataset was preserved and the resulting expression transformed to a value between 1 and 100. These transformed values could then be used to directly compare ranks of RNA expression across datasets. However, we further removed those percentile values below the 25^th^ percentile to boost high confidence signal without a substantial bias. We next used this post-percentile-coded and noise-removed matrix to carry out meta-analysis based on correlation PCA, *t*-SNE, and SC3 consensus clustering, aiming to reveal the major influencing factors that control cellular and pluripotent states.

### PCA

The percentile normalized and mRNA noised removed datasets were used to construct a Pearson correlation-based matrix (*A*, a gene expression correlation versus gene expression correlation matrix) that accounts for 100% variations of gene expression profiles. The inverse correlation matrix (*A^−1^*) was further employed to calculate the principal components (PCs, known as eigenvalues) (e.g., PC1, PC2, and PC3) by orthogonal decomposition using R programming. For example, PC1, PC2, and PC3 represent the sum of weighted correlation (*W_i_*) for each individual gene expression correlation (*G_i_*) in each sample (*S_j_*) in one column. Thus, these PC values were used to map the data points in PCA scatter plots, in which each data point contains the genome-wide gene expression correlation profile of one sample (or cell type).

### *t*-SNE visualization and cluster analysis

The *t*-SNE analysis was implemented by the R program based on a curated meta-information table, in which the details per sample can be differentiated by colored clusters in plots. For example, we can assign colors to all samples based on the names of datasets, species (e.g., mouse versus human), pluripotent states (e.g., naive, naive-like, and primed), and cellular states (e.g., undifferentiated versus differentiated). Of note, the *t*-SNE plots may differ from each other using the same datasets for analysis at a different time (van der Maaten and Hinton, 2008).

### SC3 consensus clustering

To increase the strength of the PCA and *t*-SNE analysis, we integrated SC3 consensus clustering into PCA and *t*-SNE plots. This gene expression clustering method was based on Euclidean, Pearson, and Spearman distances using the SC3 consensus clustering (http://bioconductor.org) provided by the R package (Kiselev et al., 2017).

The SC3 package includes the function named “sc3_plot_consensus” that allows us to evaluate the sample-cluster relationships for a selected number of clusters (*k*). We explored sample versus cluster assignments over a wide range of *k* using this function and found *k* = 22 to arguably be the best one, which provided a diagonal consensus matrix that was neither over-nor under-clustered.

To determine top gene markers in each cluster, we also applied the above “sc3” function to export both differentially expressed features across the samples (regardless of cluster assignments) and candidate markers per cluster. The mean cluster expression values (under all cell samples of each cluster) were used to construct a binary classifier prediction for a given gene. By default, features with an area under the receiver operating characteristic (AUROC) curve is used to quantify the accuracy of the classifier prediction of gene markers in SC3 clusters. Through the Wilcoxon signed rank test, a *P*-value was given to each gene (Supplementary Table 7). To define a gene marker, the AUROC was set to 0.8 with a 0.05 *P*-value threshold.

To better view the relationship between clustering analysis and pluripotent states, we integrated the supervised into unsupervised analyses by introducing cluster breakpoints and by aligning the clusters with cellular or pluripotent state information (Fig. 5, 7; Supplementary Table 7). The cluster fragments provide a quick view of the interchangeability of cluster rearrangements. The outcomes of gene cluster rearrangements depend on the size(s), function, redundancy, numbers of the gene cluster input.

### Web resources used in study

https://www.ncbi.nlm.nih.gov

http://bioconductor.org/packages/release/bioc/vignettes/SC3/inst/doc/SC3.html

https://distill.pub/2016/misread-tsne/

https://www.genecards.org

https://www.uniprot.org

## Supporting information

Supplementary Table 1 Curated data information

Supplementary Table 2 Datasets and protocol summary

Supplementary Table 5 post_percentile coded data

Supplementary Table 6 post_filtering pre_intersection data

Supplementary Table 7 SC3 clusters & gene markers

## Acknowledgments

This work was supported by the Intramural Research Program of the NIH at the National Institute of Neurological Disorders and Stroke. We thank Dr. Kyeyoon Park, Dr. Paul Tesar, and Dr. Pamela Robey for helpful discussions.

## Author contributions

K.R.J. and K.G.C. designed meta-analysis; K.G.C. and B.S.M. performed experiments; K.R.J. and K.G.C. prepared all figures and Supplementary Information; K.G.C. wrote the manuscript; All authors analyzed data, edited and reviewed the manuscript.

## Data availability

All raw microarray and RNA sequencing files are available online at the Gene Expression Omnibus and Sequence Read Archive (SRA) (www.ncbi.nlm.nih.gov), and the ArrayExpress Archive of Functional Genomics Data (www.ebi.ac.uk).

## Additional information

Correspondence and requests for materials should be addressed to K.R.J. or K.G.C.

---

**Supplementary Figure 1.**
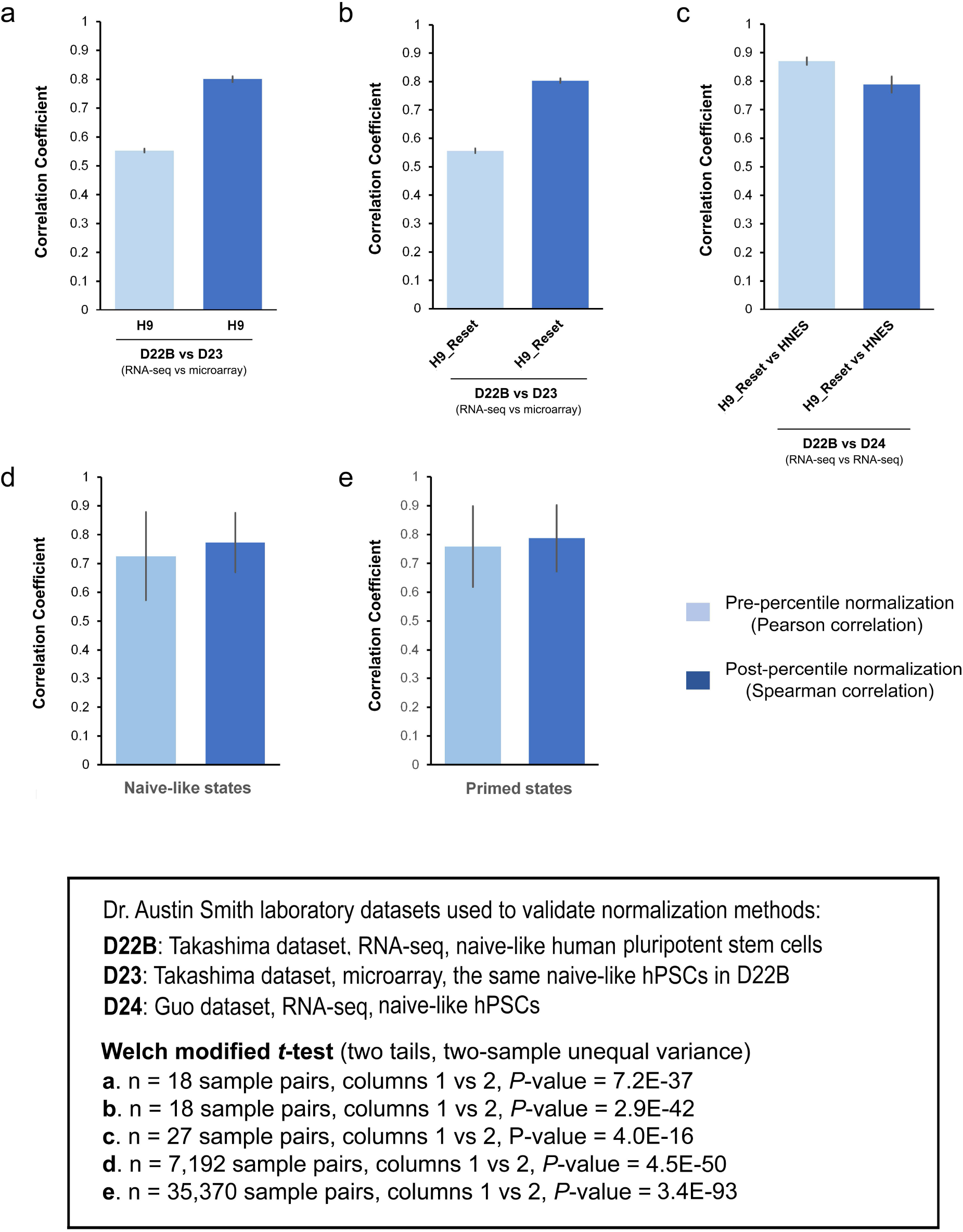

**Supplementary Figure 2.**
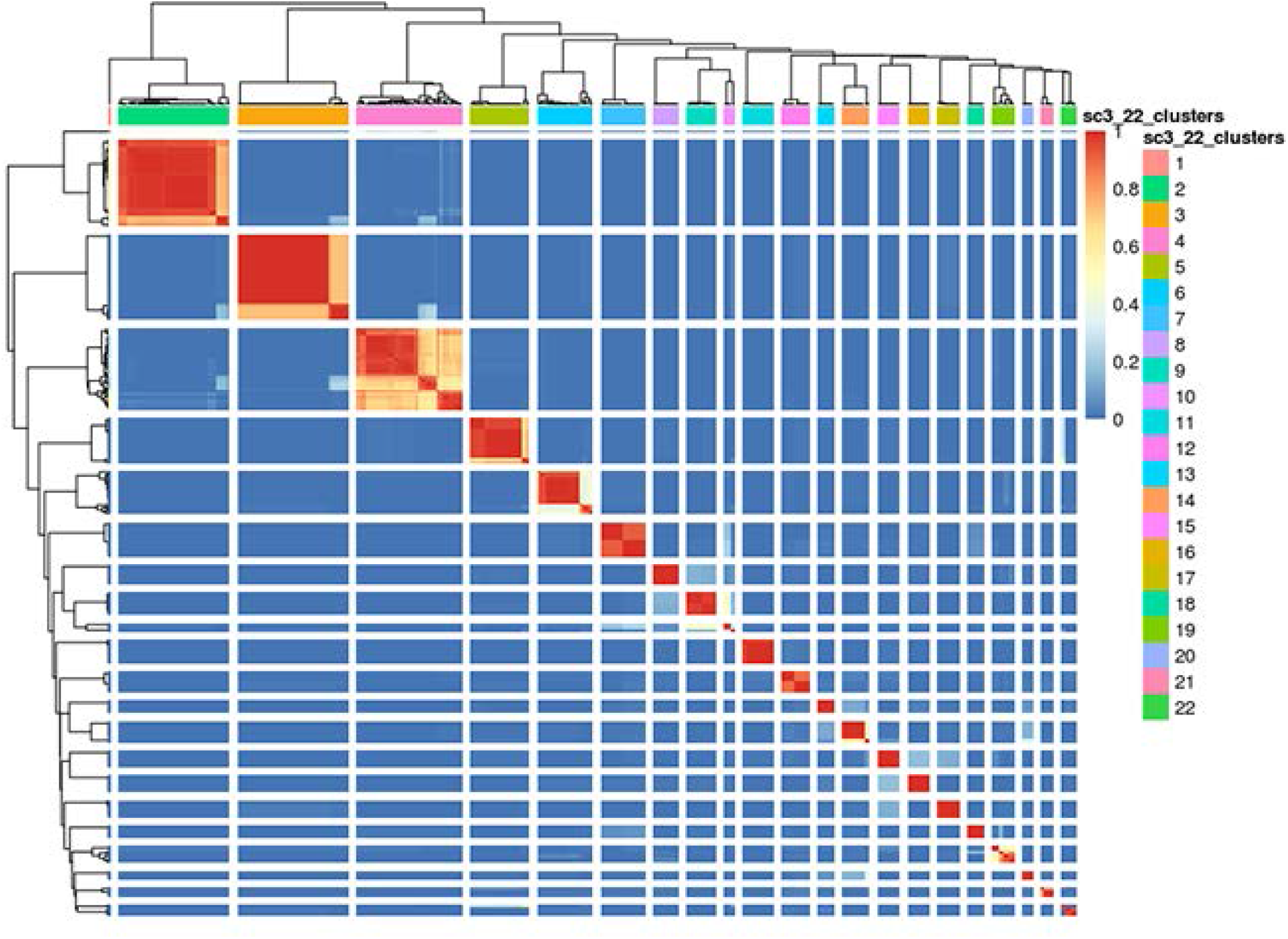

**Supplementary Table 3.** RNA processing information on data analysis. **Footnotes:** * Both microRNA (miRNA) and small nucleolar RNA (snoRNA) precursors (i.e., pri-miRNA and pre-snoRNA), which are transcribed by RNA polymerase II and possess poly(A) tails (Lee et al. EMBO J 2004, 23: 4051–60; Cai et al. RNA 2004, 10: 1957–66; Kufel and Grzechnik P. Trends Genet 2019, 35: 104-117), can be (designated as “Yes”) or cannot be (as “No” or “Less likely”) preserved (or generated) using these RNA processing and reverse-transcribing methods. **Abbreviations:** aRNA, antisense RNA; cRNA, antisense RNA; poly(A), poly(A) mRNAs; Refs, references; rRNA, ribosomal RNA; WTA2, Whole Transcriptome Amplification Kit 2. **Refs:** [1] Mallon et al. Stem Cell Res 2013, 10:57-66; [2] Chan et al. Cell Stem Cell 2013, 13: 663-675; [3] Gafni et al. Nature 2013, 504: 282-6; [4] Valamehr et al. Stem Cell Reports 2014, 2: 366-81; [5] Theunissen et al. Cell Stem Cell 2014, 15: 471-487; [6] Tesar et al. Nature 2007, 448:196-9; [7-9]; Takashima et al. Cell 2014, 158: 1254–1269; [10] Guo et al. Stem Cell Reports 2016, 6: 437-446; [11] Yan et al. Nat Struct Mol Biol 2013, 20:1131-9; [12] Vassena et al. Development 2011, 138: 3699-709.

| Dataset | Accession numbers | Expression/ Platforms | Sample RNA / cDNA preparations | Small RNAs in samples* | Potential influence on RNA detection | Refs |
| --- | --- | --- | --- | --- | --- | --- |
| D1 | GSE32923 | Microarray<br>Agilent-014693<br>GPL6480 | • Total RNAs<br>• DNA-free<br>• No rRNA depletion | • Yes | • Allow to measure small RNAs | 1 |
| D3 | E-MTAB-2031 | RNA-Seq<br>HiSeq 2000,<br>Illumina | • Poly(A) mRNA | • No<br>• Less likely | • May allow to measure small RNA precursors<br>• Defined by cluster 28, deficient in small RNAs | 2 |
| D5 | GSE46872 | Microarray<br>Affymetrix<br>Human Gene 1.0 ST | • Total RNAs<br>• DNA-free,<br>• No rRNA depletion | • Yes | • Allow to measure small RNAs | 3 |
| D6 | GSE50868 | Microarray<br>Human<br>Genome<br>U133-plus-2.0<br>chips | • Total RNAs<br>• No rRNA depletion<br>• Oligo-dT-based cDNA and aRNA | • Yes | • Allow to measure small RNAs | 4 |
| D7 | GSE59435 | Microarray<br>AFFYMETRIX<br>901837 | • Total RNAs<br>• DNA-free,<br>• No rRNA depletion<br>• Oligo-dT-based cDNA and aRNA | • Yes | • Allow to measure small RNAs | 5 |
| D18 | GSE7866 | Microarray<br>Agilent-014868<br>GPL4134 | • Total RNAs<br>• No rRNA depletion<br>• Oligo-dT-based cDNA and cRNA | • Yes | • Allow to measure small RNAs | 6 |
| D22A | E-MTAB-2857 | RNA-Seq<br>Illumina HiSeq 2000 | • Total RNAs<br>• DNA-free<br>• rRNA depletion<br>• Oligo-dT- and hexamer -based cDNA library | • Yes | • Sequencing read in 100-bp paired-end format.<br>• Allow to measure small RNA precursors | 7 |
| D22B | E-MTAB-2857 | RNA-Seq<br>Illumina HiSeq 2000 | • Total RNAs<br>• DNA-free<br>• rRNA depletion<br>• Oligo-dT- and hexamer -based cDNA library | • Yes | • Sequencing read in 100-bp paired-end format.<br>• Allow to measure small RNAs and their precursors | 8 |
| D23 | E-MTAB-2856 | Microarray<br>Affymetrix<br>Human Gene<br>Array 1.0 ST<br>arrays | • Total RNAs<br>• DNA-free<br>• Hexamer-based cRNA probe | • Yes | • Allow to measure small RNAs | 9 |
| D24 | E-MTAB-4461 | RNA-Seq<br>Illumina HiSeq 2500 | • Total RNAs<br>• DNA-free<br>• rRNA depletion<br>• Oligo-dT- and hexamer -based cDNA library | • Yes | • Sequencing read in 125-bp paired-end format<br>• Allow to measure small RNAs and their precursors | 10 |
| D25 | GSE36552 | RNA-Seq<br>Illumina HiSeq 2000 | • mRNAs<br>• rRNA depletion (<5%)<br>• cDNAs by poly(T) primer with anchor sequence (UP1) | • No<br>• Less likely | • May allow to measure small RNA precursors<br>• Defined by 6 SC3 clusters, 2/6 clusters enriched with small RNAs | 11 |
| D26 | GSE29397 | Microarray:<br>Affymetrix<br>Human Gene<br>ST 1.0 arrays | • Total RNA,<br>• DNA-free | • Yes | • Product size ranges from 100–1000 bases, typically | 12 |
|  |  |  | • primers composed of a semi-degenerate 3' end and a universal 5' end<br>• WTA2 cDNA library |  | smaller for degraded RNAs<br>• Allow to measure small RNA precursors and small RNAs |  |

**Supplementary Table 4.**
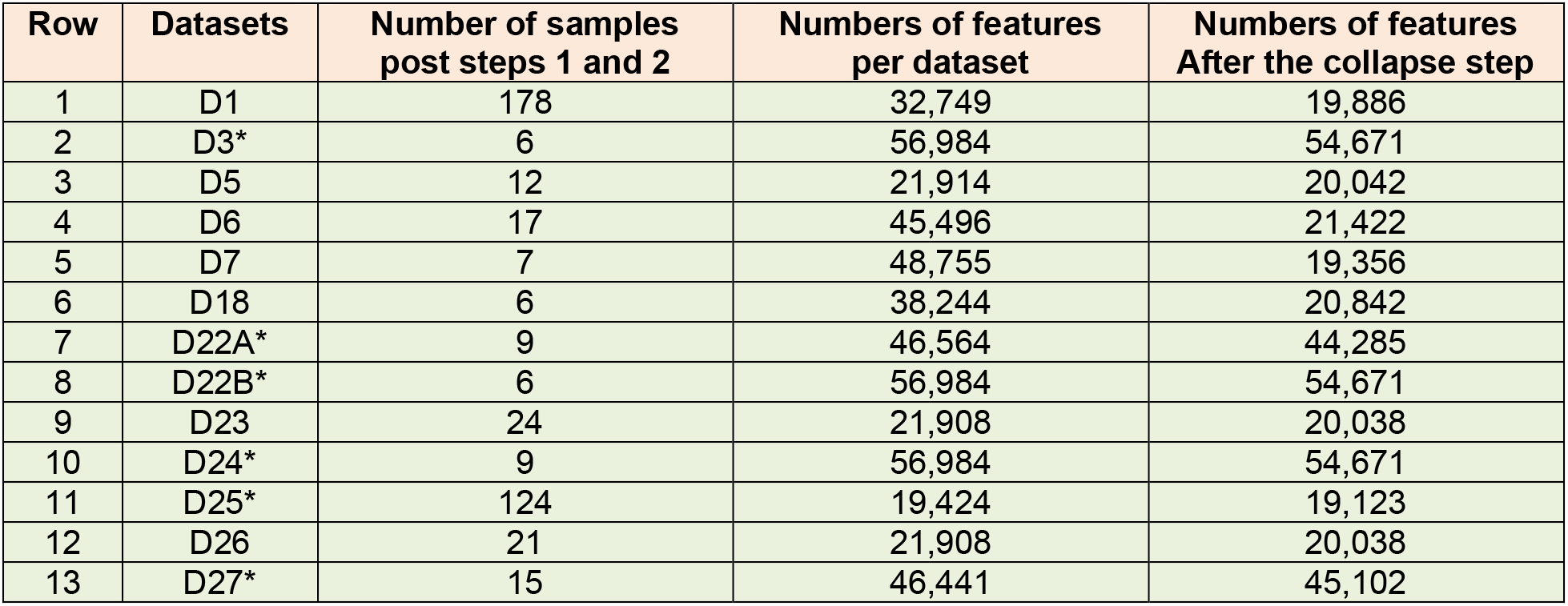
Number of features with detectable expression (i.e., entries) in at least one sample per dataset. **Footnotes:** **Colum 3:** Numbers of samples per dataset after Log2 transformation, quantile normalization, and outliner removal are shown in this column. **Column 4:** Gene features that were not assigned to an annotation were discarded. Numbers of features per data set after this step are listed in this column. **Column 5:** Gene features representing the same annotation were collapsed into single expression by taking the observed maximal values. As indicated, there is no strong bias in the number of features detected versus undetected across the RNAseq (as indicated by asterisk signs) and microarray datasets. We determined the number of gene features (symbols) that must detectable expression greater than the 25th percentile in at least one sample within a dataset. There are total 11,608 unique intersecting gene features (symbols) across the 13 datasets (including 7 microarrays and 6 RNA-seq).

